# An epigenetic memory of inflammation controls context-dependent lineage plasticity in the pancreas

**DOI:** 10.1101/2021.11.01.466807

**Authors:** David J. Falvo, Adrien Grimont, Paul Zumbo, Julie L. Yang, Alexa Osterhoudt, Grace Pan, Andre F. Rendeiro, John E. Wilkinson, Friederike Dündar, Olivier Elemento, Rhonda K. Yantiss, Doron Betel, Richard Koche, Rohit Chandwani

**Author notes:** Lead contact: Rohit Chandwani, MD, PhD; Correspondence.

## Abstract

Inflammation is essential to the disruption of tissue homeostasis, and, in the pancreas, can destabilize the identity of terminally differentiated acinar cells. Herein we employ lineage-traced mouse models to delineate the chromatin dynamics that accompany the cycle of metaplasia and regeneration following pancreatitis, and unveil the presence of an epigenetic memory of inflammation in the pancreatic acinar cell compartment. We observe that despite histologic resolution of pancreatitis, acinar cells fail to return to their molecular baseline after several months, representing an incomplete cell fate decision. *In vivo*, this epigenetic memory controls lineage plasticity, with diminished metaplasia in response to a second inflammatory insult but increased tumorigenesis with an oncogenic *Kras* mutation. We demonstrate that both persistent chromatin and transcriptional changes constituting memory are recalled with oncogenic stress. Together, our findings define the dynamics and recall of an epigenetic memory of inflammation that impacts cell fate decisions in a context-dependent manner.

## INTRODUCTION

Biological systems operate within fluctuating environments, and, therefore, are inherently tasked with accurately responding in real-time to a myriad of signals. Recent evidence suggests that a memory of inflammation can be encoded and retained in the epigenome of cells even following resolution of the initial stimulus (Ostuni et al., 2013; Naik et al., 2017; Ordovas-Montanes et al., 2018). The presence of ‘inflammatory memory’ suggests that preservation of tissue homeostasis also incorporates an evolutionary adaptation in which future responses are educated by past experiences.

In the pancreas, inflammation, which is known to play varying roles in both tumor initiation and progression, results in a reversible cell-fate transition known as acinar-to-ductal metaplasia (ADM). Homeostasis following ADM is typically restored by prompt regeneration of the acinar compartment (Means et al., 2005; Fendrich et al., 2008), given the absence of a long-lived progenitor pool in the adult pancreas (Kopp et al., 2011). By contrast, in the context of mutant *Kras*, ADM is a first step that precedes the development of pancreatic intraepithelial neoplasia (PanIN) and subsequent pancreatic ductal adenocarcinoma (PDAC). The earliest preclinical models employing oncogenic *Kras* activated by embryonic Cre drivers demonstrated sufficiency of mutant *Kras* to unveil PanIN and PDAC (Grimont et al., 2021). However, in the adult mouse, inflammation, typically experimentally elicited by the cholecystokinin analog caerulein, is necessary to give rise to neoplastic lesions that are most effectively derived from acinar cells (Guerra et al., 2007; Guerra et al., 2011; Habbe et al., 2008; Kopp et al., 2012). Recent data highlight the capacity of inflammation to destabilize pancreatic epithelial cell identity (Cobo et al., 2018; Alonso-Curbelo et al., 2021) to promote tumor initiation via acquisition of progenitor features (Means et al., 2005; Li et al., 2021) and/or outgrowth of specific critical niche populations (Westphalen et al., 2016). These models have thus demonstrated robust cooperativity between oncogenic stress and contemporaneous inflammation.

In the patient setting, it has been observed that chronic pancreatitis is a well-established risk factor for pancreatic cancer. Surprisingly, a single episode of self-limited acute pancreatitis is suggested to confer an increased risk of developing subsequent PDAC up to 10 years after the episode (Kirkegard et al., 2018). Recent evidence offers one potential explanation for this phenomenon, wherein remote inflammation can support tumorigenesis (Del Poggetto et al., 2021). However, it remains unknown what chromatin domains are associated with inflammatory memory, how memory evolves over time, and how it is recalled with secondary responses. Finally, whether inflammatory memory potentiates nascent tumors or supports initial lineage plasticity is an open question.

Here, we leverage lineage tracing of pancreatic acinar cells to define the long-term ramifications of a transient inflammatory insult on pancreatic tissue homeostasis and tumorigenesis. We find that inflammation has a prolonged and durable impact on acinar cell identity, and this manifests as an incomplete cell fate decision. The failure to fully regenerate alters the responsiveness to a secondary inflammatory insult and to delayed activation of mutant *Kras*. Importantly, we find that the persistent molecular alterations that constitute inflammatory memory are amplified in secondary responses, suggesting ‘recall’ of the initial insult. Our data thus highlight the ability for remote inflammation to become encoded as a lasting acinar cell-specific epigenetic memory of the prior insult and to affect lineage plasticity in a context-dependent manner.

## RESULTS

### A transient inflammatory episode induces persistent molecular alterations following pancreatic regeneration

We first asked whether a transient inflammatory episode persistently alters the transcriptional profile and chromatin accessibility landscape of pancreatic acinar cells. We used *Mist1*-Cre^ERT2^; LSL-tdTomato (MT) mice to conditionally restrict tdTomato expression to the acinar compartment after tamoxifen treatment (Habbe et al., 2008). After tdTomato activation, a transient inflammatory insult was induced in MT mice via intraperitoneal administration of caerulein (or saline as a control) using a 3 week protocol, followed by recovery intervals of 2 days and 3 weeks (Figure 1A). We observed that after 2 days of recovery, pancreatic histology exhibited pronounced metaplasia and tissue damage (Figure 1B). At 3 weeks of recovery, the pancreas was devoid of ADM and inflammatory infiltrates, with absence of the ductal marker CK19 from acinar cells, indicating redifferentiation of the acinar compartment (Figure 1C).

**Figure 1.**
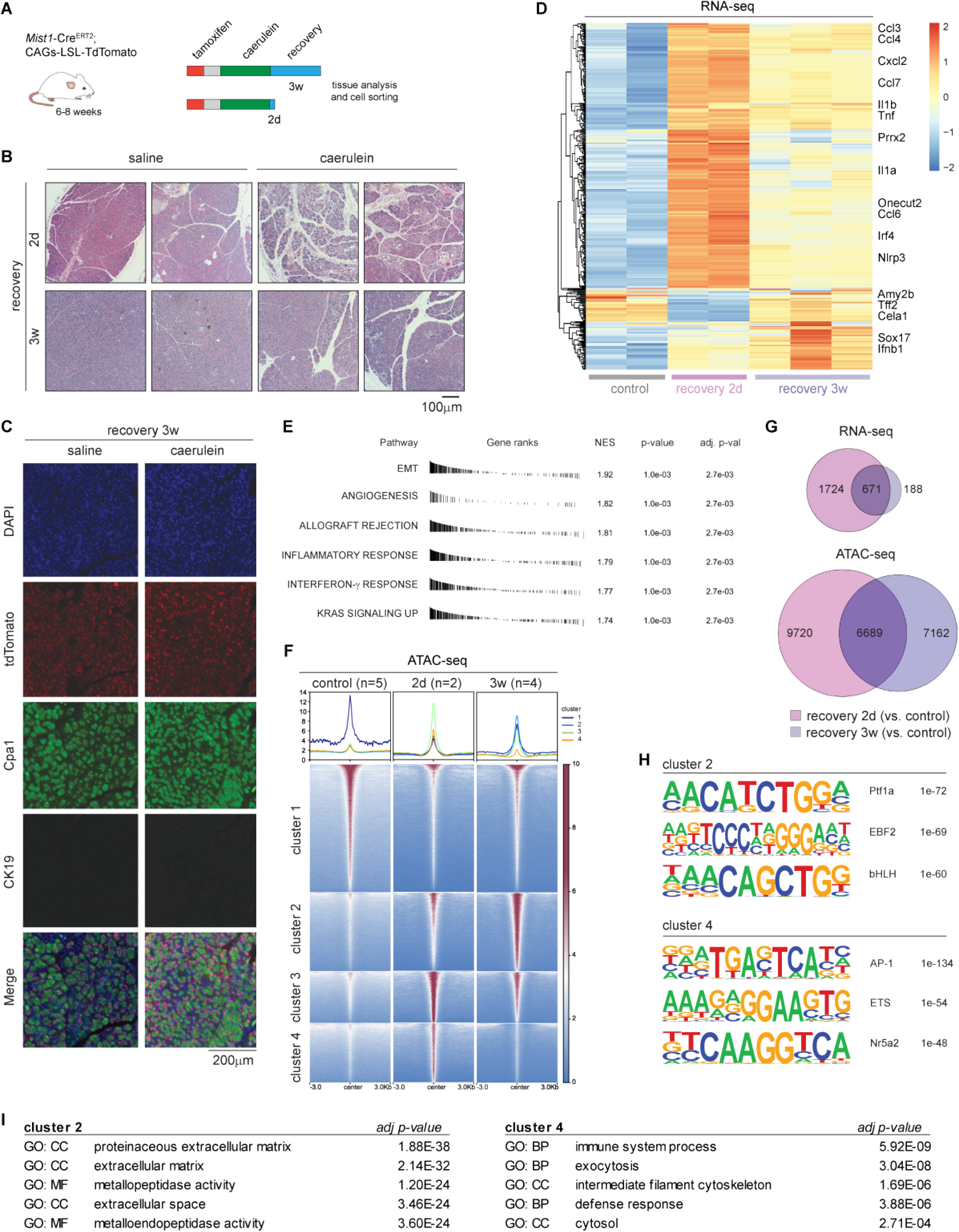
A transient inflammatory episode induces persistent molecular alterations following pancreatic regeneration. **A**, Schematic representation of lineage-traced mouse model with treatment regimen. **B**, Hematoxylin and eosin staining of mouse pancreas sections collected 2 days and 3 weeks after caerulein-induced pancreatitis. **C**, Immunofluorescence for DAPI, tdTomato, Carboxypeptidase A1 (Cpa1), and Cytokeratin 19 (CK19) staining of pancreas sections collected 3 weeks after pancreatitis. Images are representative of N = 3 mice per condition. **D**, Heatmap of RNA-seq data generated with 417 differentially expressed genes (thresholds: log_2_(fold change) > abs(2); p-value < 0.01) when comparing inflammation-naïve tdTomato(+) acinar cells (control) and peak inflammation tdTomato(+) acinar cells (recovery 2d); supervised clustering according to indicated condition. Each column is a biological replicate mouse. **E**, Gene Set Enrichment Analysis (GSEA) of RNA-seq data between control and recovery 2d condition, top six results are shown. **F**, Tornado plots visualizing the chromatin accessibility upstream and downstream of a 6-kilobase window (i.e. -3kb to +3kb) at different locations throughout the genome (the clusters and genomic locations populating each cluster are identical to the sites in Figure 1F). Red indicates maximal signal and blue signifies absent signal. Each row corresponds to a genomic location, and each column represents the average signal across N=2-5 biological replicate mice treated according to the conditions indicated. **G**, Venn diagrams illustrating the degree of overlap between 2 days of recovery versus control and 3 weeks of recovery versus control from a transcriptional (top) and chromatin accessibility (bottom) standpoint; log2FC>1; FDR<0.05. **H**, HOMER analysis depicting top three motif enrichments in Cluster 2 and Cluster 4 and associated adjusted p-values. **I**, GO term enrichment for genes associated with differentially accessible sites in Cluster 2 and Cluster 4. Top five results are shown along with adjusted p-values. CC = cellar component; BP = biological process; MF = molecular function.

To assess the spectrum of molecular changes induced during and after inflammation, pancreata were harvested from both MT mice treated with caerulein, at 2 days and 3 weeks after the insult. tdTomato(+) acinar cells were isolated via fluorescence-activated cell sorting (FACS) for downstream profiling of transcriptional and chromatin accessibility changes using RNA-seq and ATAC-seq, respectively. At 2 days after injury (recovery 2d), we observed an expected downregulation of acinar-specific genes (e.g. *Amy2b, Tff2, Cela1*), with strong upregulation of ductal (e.g. *Krt19, Onecut2)* and pro-inflammatory genes (e.g. *Ccl3, Ccl4, Cxcl2, Il1b, Tnf*). Surprisingly, we found that after 3 weeks (recovery 3w), acinar cells display a mixed transcriptional state, characterized by persistent expression of metaplasia-associated genes despite normal histology (Figure 1D). Using Gene Set Enrichment Analysis (GSEA), we observed an enrichment of transcripts involved in epithelial-to-mesenchymal transition (EMT), angiogenesis, inflammation, and related to Kras signaling at 3 weeks of recovery –- albeit to a lesser extent than at 2 days – suggesting incomplete regeneration at this timepoint (Figure 1E; Figure S1A).

Next, we evaluated the changes in chromatin accessibility at the peak of inflammation and after resolution of pancreatitis. Principal Component Analysis (PCA) on the ATAC-seq data from FACS-sorted acinar cells collected from all three conditions showed separation between recovery 2d and recovery 3w mice; consistent with the RNA-seq data, chromatin accessibility remained altered from baseline after 3 weeks of recovery (Figure S1B). Inflammation-exposed tdTomato(+) acinar cells collected after 2 days of recovery exhibited dramatically reduced accessibility at genomic regions enriched in control acinar cells (Figure 1F; Cluster 1), suggesting a loss of acinar cell identity. Additionally, we observed profound gains in accessibility at intergenic and intronic regions unveiled during peak inflammation (Figure 1F; Clusters 3 and 4). Interestingly, we found that there were novel gains of accessibility at 3 weeks of recovery (Figure 1F; Cluster 2) and dramatic losses of accessibility in Cluster 4, highlighting ongoing molecular changes in the acinar compartment despite evidence of regeneration by histology.

To understand the scope of these molecular alterations, we compared the degree of overlap between 2 days and 3 weeks of recovery for both the transcriptional and chromatin accessibility changes observed. Whereas the majority of persistent transcriptional changes at 3 weeks were a subset of the ‘recovery 2d’ differentially expressed genes (DEGs), the differentially-accessible regions (DARs) specific to 3 weeks of recovery were greater in number and more distinct (Figure 1G). This suggests that chromatin – to a greater extent than the transcriptome – remains dynamic up to 3 weeks after a transient insult.

Next, we performed motif analysis on the clustered DARs, identifying a number of acinar-specific factors (e.g. Ptf1a, Bhlha15) in Cluster 2, while motifs for factors involved in pancreatitis were found in Cluster 4 (e.g. AP-1, Nr5a2) (Figure 1H; Figure S1C). Broadly, 3 weeks of recovery restored accessibility of regions enriched in motifs for lineage-specifying TFs (Nr5a2, Bhlha15, Nkx6.1) but showed persistent depression of AP-1 motifs (Figure S1D). Gene Ontology (GO) enrichment analysis on the clustered DARs identified a number of processes associated with the extracellular matrix (ECM) in Cluster 2 (Figure 1I), suggesting that there are ongoing changes in tissue architecture at 3 weeks of recovery, perhaps in keeping with the loss of cellularity seen by H&E (Figure 1B). In Cluster 4, where there is rapid resolution of transient increases in accessibility, we found GO terms associated with processes characteristic of pancreatitis (e.g. immune system process, exocytosis). These findings demonstrate that inflammation-resolved acinar cells retain expression of genes upregulated during pancreatitis. However, these transcriptional changes are relatively small compared to the chromatin changes, where acinar cells retain and even accumulate clear gains in accessibility at sites normally absent in control acinar cells, while also failing to re-establish baseline accessibility at acinar-specific locations.

### Memory of inflammation manifests as an incomplete cell fate decision

Because we observed emergent acinar cell alterations at 3 weeks following pancreatitis, we asked how these molecular dynamics evolve over time. We therefore treated mice as previously described, but extended the recovery window to include 6, 12, and 18-week recovery timepoints. At 12 and 18 weeks of recovery, we saw continued evidence of normal tissue architecture (Figure 2A), and no persistent changes to the abundance of the ADM/ductal markers Sox9 and CK19 (Figure S2A-B) nor the acinar marker Cpa1 (Figure S2B). We evaluated the persistence of phospho-Erk (Indicating MAPK pathway activation), as well as the progenitor markers Klf5 and Nestin; while these were robustly induced at 2 days, there was no lasting expression at 12 weeks (Figure 2B).

**Figure 2.**
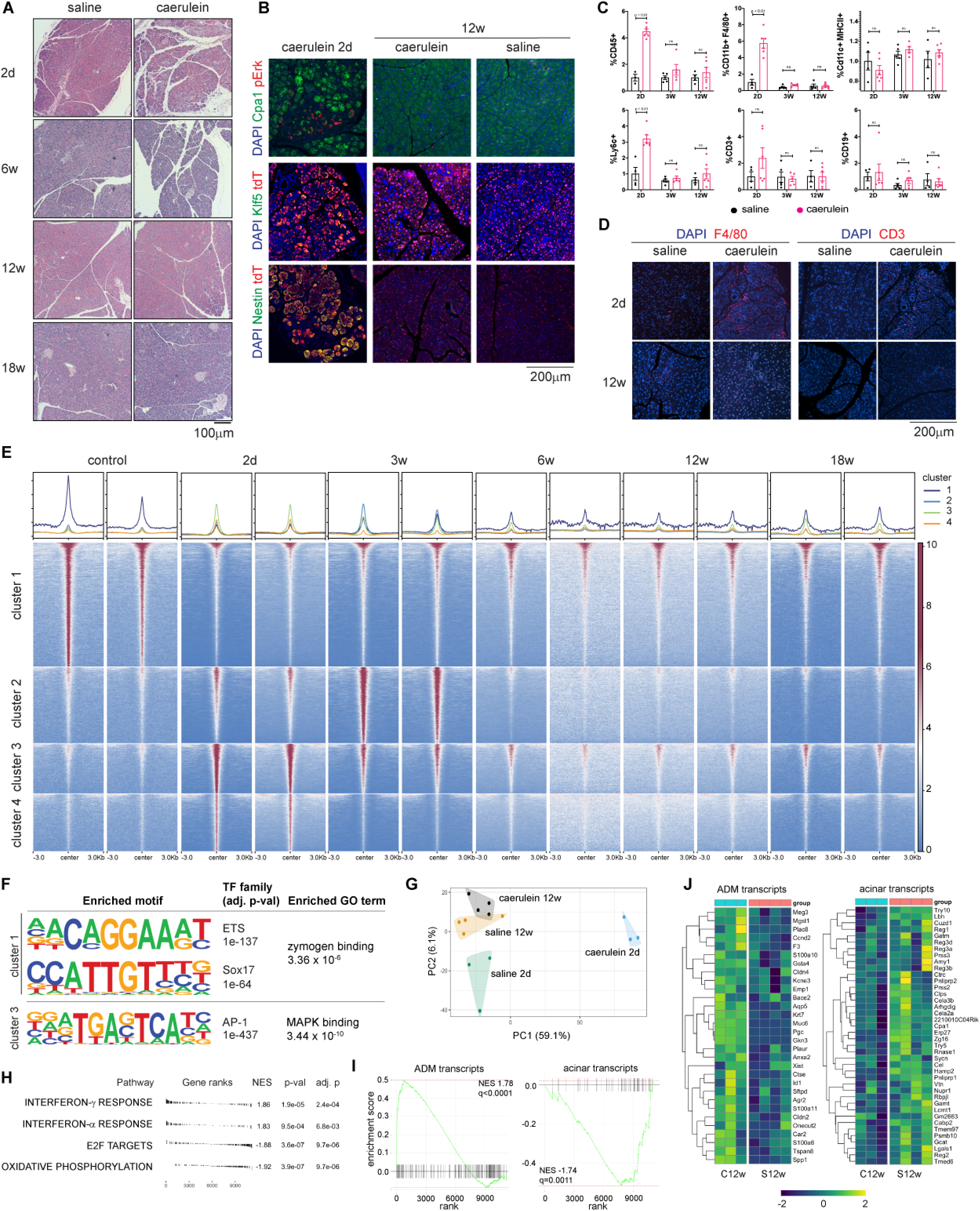
Memory of inflammation manifests as an incomplete cell fate decision. **A**, Hematoxylin and eosin staining of naïve (saline) or prior inflammation (caerulein-treated) mouse pancreas sections collected after 2 days, 6, 12, and 18 weeks of recovery. **B**, Immunofluorescence for Cpa1, tdTomato, phosphor-Erk, Klf5, Nestin and DAPI staining of pancreas sections collected 2 days and 12 weeks after pancreatitis (or control). Representative images shown are from a total of N=2-5 mice per condition. **C**, Flow cytometric quantification of CD45(+), CD11b(+) F4/80(+), CD11c(+) MHCII(+), Ly6c(+), CD3(+), CD19(+) immune cell populations showed in percentage of isolated cells from pancreata of inflammation-naïve (saline) and inflammation-exposed (caerulein) mice collected after 2 days, 3 weeks, and 12 weeks of recovery. Student’s t-test was performed between conditions. N=3-5 mice per condition. **D**, Immunofluorescence for F4/80, CD3 and DAPI of pancreas sections collected 2 days and 12 weeks after pancreatitis. **E**, Tornado plots visualizing the chromatin accessibility upstream and downstream of a 6-kilobase window (i.e. -3kb to +3kb) at different locations throughout the genome (the clusters and genomic locations populating each cluster are identical to the sites in Figure 1F). Red indicates maximal signal and blue signifies absent signal. Each row corresponds to a genomic location, and each column represents a single biological replicate mouse treated according to the conditions indicated. Two mice per condition are shown, representative of N=2-5 mice per condition. **F**, HOMER analysis depicting motif enrichment in Cluster 1 and Cluster 3 (with adjusted p-values), as well as the top GO term (with associated adjusted p-value) associated with sites enriched Cluster 1 and Cluster 3. **G**, Principal Component Analysis (PCA) plot of bulk RNA-seq data generated from tdTomato(+) acinar cells collected 2 days and 12 weeks after either pancreatitis or control treatment. **H**, GSEA of RNA-seq data derived from inflammation-naïve tdTomato(+) acinar cells versus inflammation-exposed tdTomato(+) acinar cells (12 weeks of recovery). Top 4 most up-or down-regulated pathways are shown. **I**, GSEA enrichment plot of RNA-seq data derived from prior inflammation (caerulein + 12w recovery) vs. naïve (saline + 12w recovery) samples for the indicated ADM and acinar gene sets (obtained from ref. 16) **J**, Heatmap of RNA-seq data delineating the leading edge ADM and acinar transcripts in C12w and S12w samples; each column represents a single biological replicate mouse.

We then asked if there was a persistent microenvironment change that accompanied the recovery from pancreatitis. Profiling of the immune compartment via flow cytometry showed robust infiltrates of macrophages and monocytes at 2 days that returned to baseline levels with 3 and 12 weeks of recovery. No accumulation of dendritic cells, T cells, or B cells was evident (Figure 2C). By immunofluorescence, we also saw no evidence of residual infiltration of macrophages and T cells by F4/80 and CD3 staining, respectively (Figure 2D). To confirm there were no stable perturbations to cell type composition after pancreatitis, we also performed single-cell RNA-sequencing on the entire pancreas using a 2-day protocol to potentiate the severity of acute inflammation. This showed neither a persistent immune infiltrate at 6 weeks, nor a shift in transcriptional state among immune cells, fibroblasts, or endothelial cells (Figure S2C). Importantly, we did detect the abundant infiltration of myeloid cells and emergence of ADM cells at 2 days, but no persistent alteration over time, recapitulating what we observed using the 3 week caerulein protocol (Figure 2C-D). In the epithelial compartment, we observed that ADM cells at 2 days manifest as a mixed transcriptional state, characterized by the expression of genes enriched in both acinar and ductal cells (Figures S2D and E); however, we noted no persistent alterations by scRNA-seq over time.

Our data therefore suggested a restoration of tissue architecture, gene expression, and microenvironment composition with prolonged recovery. Next, we determined if chromatin alterations in acinar cells persist despite these findings. To this end, we obtained sorted tdTomato(+) acinar cells from the 6, 12, and 18-week recovery timepoints for subsequent downstream profiling of chromatin accessibility. Unexpectedly, we found that even after 18 weeks of recovery, acinar cells retained a chromatin signature of prior inflammation (Figure 2E; Cluster 3). Motif enrichment of these gained memory regions showed an abundance of AP-1 motifs, with a top gene ontology (GO) term corresponding to MAPK pathway (Figure 2F). Conversely, acinar cell identity regions in Cluster 1 failed to completely restore chromatin accessibility enriched in controls, even after 12 and 18 weeks of recovery. We found that the sites in Cluster 1 were enriched for ETS factor and Sox17 motifs, known to be involved in pancreas development, and identified GO terms associated with processes important for acinar cell function (i.e. zymogen binding) (Figure 2F). Interestingly, regions enriched in Cluster 2 that initially increased accessibility up to 3 weeks of recovery were markedly lost at 6 weeks of recovery and beyond, suggesting that the full range of chromatin dynamics are more adequately revealed with prolonged recovery (Figure 2E). Together, these data highlight a prolonged cell-intrinsic effect of prior inflammation on chromatin states that is not manifested histologically.

To determine if any persistent transcriptional changes accompanied the chromatin dynamics, we also performed bulk RNA-seq in order to specifically interrogate sorted tdTomato(+) acinar cells in greater depth than could be achieved by scRNA-seq of the entire pancreas. We found a subtle shift in clustering of caerulein + 12 week recovery (‘prior inflammation’) samples versus saline + 12 week recovery (‘naïve’) samples (Figure 2G). No statistically significant individual gene expression changes were found at this timepoint (Figure S2F), but at the pathway level, there were increases in both inflammatory and metaplasia-specific (ADM) transcripts in prior inflammation samples; conversely, acinar transcripts (Schlesinger et al., 2020) were enriched in inflammation-naïve acinar cells (Figures 2H-J). Further, leading-edge ADM genes increased in expression in prior inflammation displayed nearby regulatory elements with persistently increased accessibility (i.e. cluster 3-type), while acinar gene-related enhancers exhibited cluster 1 dynamics (Figure S2G). These results suggest that injury establishes a memory of inflammation that manifests as an incomplete cell fate decision, where both chromatin alterations and transcriptional states show residual subtle evidence of the remote metaplastic event.

### Prior inflammation alters the capacity for subsequent metaplasia

Given these findings, we asked whether the memory of inflammation impacts acinar cell responses to future stimuli. We first tested if memory predisposes acinar cells to undergo promiscuous metaplasia. To test this, we treated wild-type C57BL/6 mice to primary inflammation (or control) followed by a recovery period and then secondary treatment with caerulein. With a recovery period of 3 weeks, we observed that inflammation elicited more robust ADM in mice recovered from prior inflammation (Figure S3A-B) suggesting that transcriptional and chromatin changes early in recovery could potentiate the impact of a secondary inflammatory insult.

We subsequently focused on allowing for 12 weeks of recovery between primary and secondary insults to better articulate the long-term consequences of prior inflammation when a steady-state appears to have been reached (Figure 3A). Surprisingly, we found pancreata exposed to prior inflammation are refractory to secondary acute inflammatory rechallenge, with increased ADM in inflammation-naïve mice compared to inflammation-resolved mice (Figures 3B-C). Co-staining of Cpa1 and CK19 was observed in inflammation-resolved tissue re-challenged with caerulein, indicative of early ADM, while Cpa1 was lost in the metaplastic lesions of inflammation naïve tissue similarly re-challenged, suggesting more advanced ADM (Figure 3D). In addition, cleaved-caspase 3 (CC3) staining co-localized with advanced metaplastic lesions, was proportional to the degree of metaplasia, and not otherwise differentially enriched depending on the prior insult (Figure 3E). We performed similar *in vivo* re-challenge experiments with fewer injections at the time of secondary caerulein, and again observed more metaplasia / transdifferentiation in inflammation-naïve pancreata as compared to inflammation-exposed pancreata (Figure S3C-D). Therefore, prior inflammation was associated with diminished lineage plasticity to further inflammatory insults.

**Figure 3.**
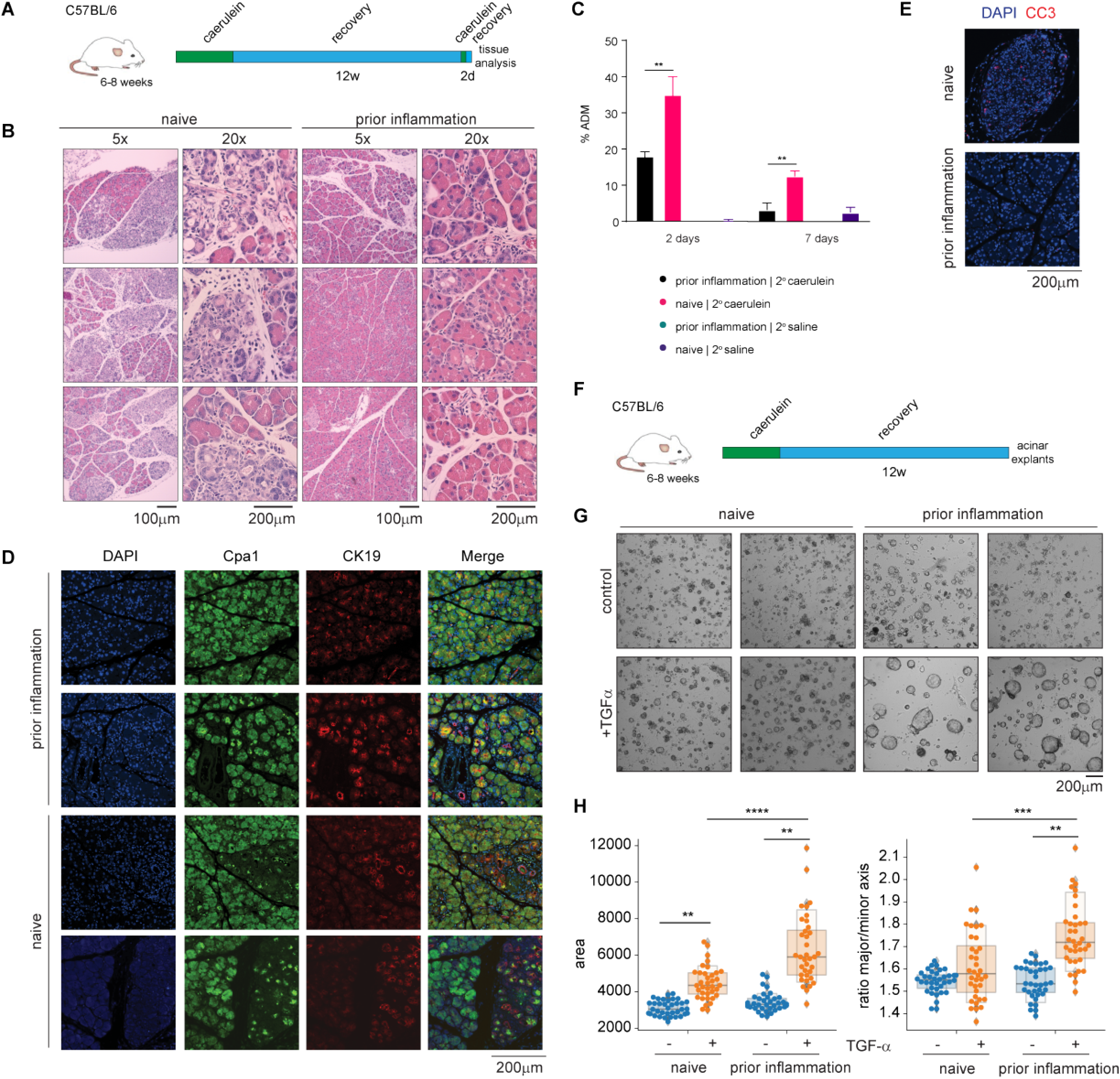
Prior inflammation alters the capacity for subsequent metaplasia. **A**, Schematic representation of primary pancreatitis and inflammatory re challenge treatment regimen in wild-type mice. **B**, Hematoxylin and eosin staining of inflammation-naYve (N = 4) and inflammation-resolved (N = 5) mouse pancreas sections collected 2 days after inflammatory re-challenge. **C**, Quantification of the ADM area (in percentage) of pancreas sections collected from inflammation-na’ive and inflammation-resolved mice re-challenged with either saline or caerulein (n=3-5 per condition). Student’s t-test was performed to compare conditions. **D**, lmmunofluorescence for Cpa1 and CK19 and DAPI staining of inflammation-na’^i^ve and inflammation-resolved mouse pancreas sections collected 2 days after inflammatory re-challenge. Representative images shown are from a total of N=2-5 mice per condition. **E**, Representative immunofluorescence for cleaved caspase 3 (CC3) and DAPI staining of inflammation-na’ive and inflammation-resolved mouse pancreas sections collected 2 days after inflammatory re-challenge. Representative images shown are from a total of N=2-5 mice per condition. **F**, Schematic representation of primary pancreatitis followed by prolonged recovery and acinar explant generation. **G**, Representative images of acinar explants generated from inflammation-na’ive and inflammation-resolved mice re-challenged *in vitro* with either vehicle or recombinant human TGFα; N=4 mice per condition. **(H)** Boxplots of image analysis quantifying the area and major/minor axis ratio for inflammation-na’ive and inflammation-resolved explants treated with vehicle or recombinant human TGFα. Each point represents an object in one of 3 wells per mouse, N=4 mice per condition. Mann Whitney was performed to compare conditions.

To further understand the phenotypic consequences of inflammatory memory, we tested if prior inflammation alters the capacity of acinar cells to transdifferentiate *in vitro* in a cell-autonomous fashion. We exposed wild-type C57BL/6 mice to primary inflammation (or control) followed by a recovery period of 12 weeks, and isolated acinar cells for embedding in Matrigel (Figure 3F). In the absence of rhTGF α, we observed no change in acinar cells from naïve or prior inflammation mice (Figure 3G, upper panels). Treatment with rhTGF α *in vitro*, which activates signaling downstream of EGFR, gave rise to ductal cystic structures (Figure 3G, lower panels). Unexpectedly, inflammation-resolved acinar cells generated substantially larger cystic structures when compared to inflammation-naïve acinar cells (Figure 3H). Our results thus highlight that the impact of prior inflammation is to diminish metaplastic lesions *in vivo*, but that EGFR-MAPK signaling pathway activation remains intact in acinar cells.

### Prior inflammation lowers the threshold for subsequent Kras-driven tumor initiation

We reasoned that the heightened response to a specific EGFR ligand would imply that prior exposure to inflammation lowers the threshold for malignant transformation. Most studies of adult mutant *Kras* activation utilize a contemporaneous inflammatory insult; therefore, we first sought to determine if any temporal separation between inflammation and oncogene activation could drive tumorigenesis. To test this, we treated MT and *Mist1*-Cre^ERT2^; LSL-Kras^G12D/+^; LSL-tdTomato (MKT) mice with either saline or caerulein as before, followed by recovery, and then activation of Kras^G12D^ with tamoxifen (Figure 4A; Figure S4A). Importantly, this system provided complete temporal control over endogenous levels of *Kras* expression and with specific restriction to pancreatic acinar lineage. With 3 weeks of recovery between injury and *Kras*^G12D^ activation (Figure S4A), we indeed observed substantial PanIN development in mice recovered from prior inflammation, as compared to rare lesions in naïve mice (Figure S4B). Extension of the temporal separation to 12 weeks (Figure 4A) showed increased ADM and PanIN in prior inflammation mice (Figures 4B, 4C [right panel], 4D). Immunofluorescent staining of both inflammation-naïve and inflammation-exposed MKT pancreas sections revealed the loss of Cpa1, as well as the colocalization of CK19, at these PanIN lesions (Figure 4E). We also found that the PanIN lesions were enriched with Dclk1^+^ cells, as previously described (Westphalen et al., 2016), but not to different degrees between comparable lesions in the two conditions (Figure 4F).

**Figure 4.**
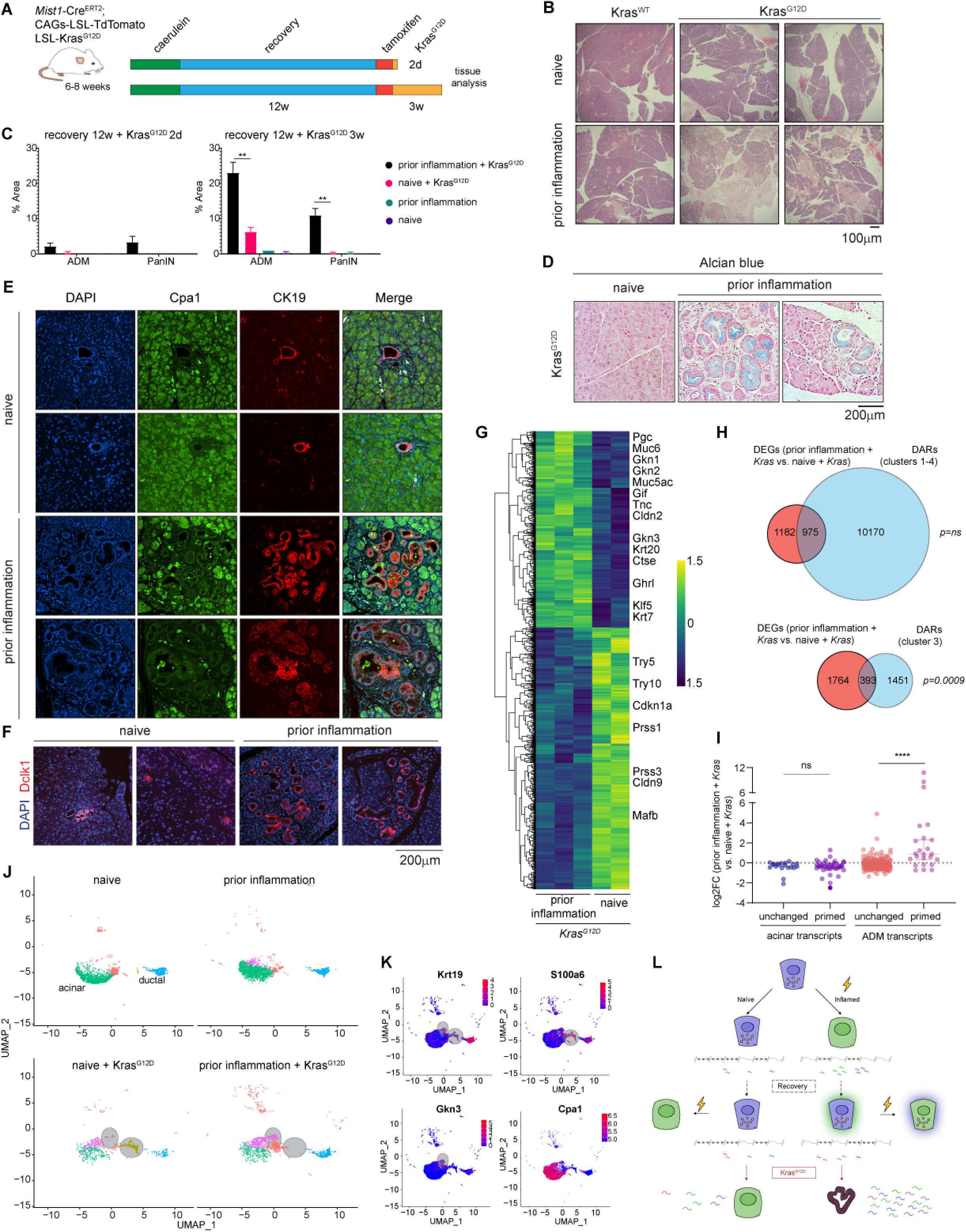
Prior inflammation lowers the threshold for subsequent Kras-driven tumor initiation. **A**, Schematic representation of lineage-traced mouse model with initial exposure to pancreatitis, prolonged recovery, and delayed Kras^G12D^ activation. **B**, Hematoxylin and eosin staining of inflammation-naïve and inflammation-exposed mouse pancreas sections collected after 12 weeks of recovery and 3 weeks of mutant Kras activation. Representative images shown are from a total of N=4-5 mice per condition. **C**, Histologic quantification of ADM and PanIN lesions in percentage of pancreas sections collected from mice exposed to the corresponding conditions wherein duration of Kras^G12D^ activation is either 2 days or 3 weeks. **D**, Alcian blue staining of pancreas sections collected from inflammation-naïve and inflammation-resolved mice exposed to 12 weeks of recovery and then 3 weeks of Kras^G12D^. Images are representative of N=4-5 mice per condition. **E & F**, Immunofluorescence for **(E)** Cpa1 and CK19 and **(F)** Dclk1 and DAPI staining of pancreas sections collected from inflammation-naïve and inflammation-resolved mice exposed to 12 weeks of recovery and then 3 weeks of mutant Kras. Representative images shown are from a total of N=2-5 mice per condition. **G**, Heatmap of differentially expressed genes between inflammation-naïve and inflammation-resolved tdTomato(+) acinar cells after 12 weeks of recovery and then 2 days of Kras^G12D^. Threshold of p<0.01 was used to identify DEGs. Selected genes are highlighted from both cluster. **H**, Venn diagrams illustrating the degree of overlap between differentially expressed genes (DEGs) altered with prior caerulein plus brief Kras^G12D^ activation and genes associated with differentially accessible regions (DARs) enriched in respective clusters. Fisher’s exact test was used. **I**, Dot plot illustrating the fold change of leading edge ‘acinar transcripts’ (primed) and leading edge ‘ADM transcripts’ (primed) compared to unchanged ‘acinar transcripts’ and unchanged ‘ADM transcripts’ (identified from Figure 2I-J) in inflammation-naïve and inflammation-resolved tdTomato(+) acinar cells exposed to 12 weeks of recovery and then 2 days of Kras^G12D^. Student’s t-test was performed to compare conditions. **J**, UMAP of single-cell RNA-seq data plotting only the epithelial cell compartment (e.g. acinar, ductal) in inflammation-naïve and inflammation-resolved tdTomato(+) acinar cells exposed to 12 weeks of recovery before (upper panels) and after (lower panels) 2 days of Kras^G12D^; gray circles highlight condition-specific cell populations. **K**, Feature plots generated with single-cell RNA-seq data illustrating the expression of ductal/PanIN and acinar markers.

To evaluate the synergistic limits of inflammatory memory combined with oncogenic Kras we then restricted the duration of Kras^G12D^ activation to a brief window of 2 days (Figure 4A). Surprisingly, even short-term activation of Kras^G12D^ was sufficient to induce neoplastic transformation in inflammation-exposed acinar cells; pancreata of inflammation-naïve mice were devoid of any PanIN, and displayed only sparse early metaplastic lesions (Figure 4C; left panel and Figure S4C). Indeed, CK19+ Cpa1-lesions (Figure S4D) and Dclk1+ lesions (Figure S4E) were found only in the context of prior inflammation. These findings indicate differential responses to oncogenic *Kras* that are driven by a temporally remote prior inflammatory episode.

To understand the molecular underpinnings of this phenotype, we performed RNA-seq on inflammation-naïve and -exposed tdTomato(+) acinar cells subjected to brief *Kras*^G12D^ activation. As expected, we found that transcripts associated with PanIN and proliferation were specifically upregulated with prior exposure to inflammation despite both cohorts having the same duration of *Kras* activation (Figure 4G; Figure S4F). We then asked whether brief *Kras*^G12D^ activation-induced genes associated with persistently accessible memory regions. Indeed, we found that DEGs between prior inflammation and naive conditions (with superimposed *Kras*^G12D^) were more likely to be associated with cluster 3 (memory) DARs than any other cluster (Figure 4H). Additionally, we found that the leading-edge ADM transcripts that define the transcriptional component of memory (see Figure 2I; ‘ADM primed’) were more likely to be differentially expressed between prior inflammation and naive mice in the context of brief *Kras*^G12D^ activation (Figure 4I). Unexpectedly, the leading-edge acinar transcripts slightly diminished in samples exposed to prior inflammation (see Figure 2I; ‘acinar primed’) were not further downregulated upon brief activation of Kras^G12D^ (Figure 4I). Among the most differentially expressed genes between prior inflammation + *Kras*^G12D^ and naive + *Kras*^G12D^ conditions were the PanIN transcripts *Pgc, Krt7, Gkn2*, and *Gkn3*, all of which have nearby putative enhancers displaying increased accessibility long after prior inflammation (i.e. cluster 3-type) (Figure S4G). These findings suggest that the memory of prior inflammation is recalled in the response to delayed *Kras*^G12D^ activation, with both primed transcripts and chromatin unveiled with oncogene activation.

To further validate these findings, we performed scRNA-seq in mice treated with saline or caerulein followed by 12w recovery, before and after 2 days of mutant *Kras* activation. In these experiments, we observed little to no differences in gene expression prior to superimposed *Kras* (Figure 4J). However, after *Kras* activation, we saw different emergent cell populations closely related to normal acinar cells. Incipient PanIN cells in the prior inflammation + *Kras*^G12D^ condition were notable for higher expression of *Gkn3* and *Krt19* and loss of *Cpa1*; by contrast, naive + *Kras*^G12D^ mice featured distinct cells resembling ADM, with high levels of *Krt19* and *S100a6* but no *Gkn3* (Figure 4K). This is in line with the mouse histology, which was notable for ADM but no PanIN in the absence of memory of prior inflammation. Together, these single-cell data confirm the presence of a divergent response to mutant *Kras* between naïve and inflammation-resolved mice that appears closely related to the molecular alterations immediately preceding oncogene activation (Figure 4L).

## DISCUSSION

To maintain tissue homeostasis, resident cells must engage in a vast array of choreographed molecular adaptations to an array of unforeseen events. We have recently learned that these adaptations, often encoded as an epigenetic memory, in turn affect how tissues respond over time to secondary insults (Ostuni et al., 2013; Naik et al., 2017; Ordovas-Montanes et al., 2018; Del Poggetto et al., 2021). Most frequently, epigenetic memory has been associated with the response to inflammation. In the pancreas, inflammation drives the infiltration of macrophages (Liou et al., 2013; Zhang et al., 2017), fibroblasts (Liu et al., 2016), and T cell subsets (McAllister et al., 2014), that together control cell-intrinsic alterations in acinar cell fate wherein they acquire duct-like features (Reichert et al., 2013) and dedifferentiated programs (Means et al., 2005; Li et al., 2021). However, in the absence of mutant *Kras*, acinar-ductal metaplasia has long been understood to be followed by regeneration of the terminally differentiated acinar compartment. Here we demonstrate that this regeneration is incomplete. Specifically, inflammation leads to the generation of a durable lineage-specific epigenetic memory in the pancreatic acinar cell. We find this memory is molecularly encoded as an incomplete regeneration following acinar-ductal metaplasia and is recalled in divergent cell fate decisions following re-challenge with severe inflammation or oncogenic stress.

Similar findings have recently been reported (Del Poggetto et al., 2021), wherein delayed activation of mutant *Kras* (using the doxycycline-inducible iKras model) in pancreatic epithelial cells was shown to lead to shorter survival in mice recovered from a transient inflammatory insult. In epithelial cells, the authors find broad increases in chromatin accessibility 28 days after acute pancreatitis.

In keeping with these prior findings, we observe that resolution of inflammation in the adult mouse pancreas occurs in the first few weeks after injury, with normalization of cell-type specific marker abundance, loss of immune cell infiltration, and restoration of tissue composition and architecture. However, we surprisingly observe that the normal histologic appearance present a few weeks after injury belies ongoing changes to both gene expression and chromatin that continue months after supposed resolution of inflammation. By extending our molecular analyses out to 18 weeks after pancreatitis, we find evidence of <u>time-dependent resolution</u> of initial broad gains in chromatin accessibility and near-complete resolution of transcriptional changes. We show that (1) chromatin defining acinar identity is acutely lost, and regained slowly over time, which we now term ‘repressed memory’; (2) specific regions are only opened briefly at the peak of inflammation; (3) regeneration features a distinct set of regulatory elements that are unveiled specifically in the first few weeks after inflammation; and (4) a ‘retained memory’ of inflammation is not a gradual accumulation of chromatin accessibility as previously described, but instead a failure to resolve completely the changes that occurred in the initial inflammation. This added complexity is a critical addition to our understanding of what epigenetic memory is, as it provides concrete insights into the modular encoding, heterogeneity, and evolution of chromatin over time.

Recent work providing evidence of inflammatory memory has either been restricted to the stem cell compartment or not specific to a single terminally differentiated lineage. Here we demonstrate by tracing the adult acinar cell that an epigenetic memory of inflammation is actually an incomplete cell fate decision, wherein ADM chromatin features are partially retained and certain acinar features are lost. Our data suggest this property is specific to the acinar compartment, whereby we observe neither lasting alterations in other cellular compartments of the microenvironment nor to cell-intrinsic expression of typical markers of cell identity in the pancreas. The cell-type specificity in our studies clarifies that acinar cell regeneration is incomplete as much as 3 months removed from the initial insult, raising the question of whether physiologic and *in vitro* differentiation and dedifferentiation also features persistence of prior cell identity. Indeed, findings in other tissues suggest that terminally differentiated cells in the colon or stem cells in the epidermis retain a memory of their embryonic (Jadhav et al., 2019) or niche (Gonzales et al., 2021) origins, respectively. Together, our data provide concrete evidence that transient cell fate decisions also can be encoded as an epigenetic memory.

Prior studies support a role for inflammatory memory in driving subsequent responses. Here we find evidence of effects on subsequent acinar cell responses, but in a time- and context-dependent manner. We find that with a short recovery window (3 weeks) the propensity for ADM after secondary inflammation is heightened by memory of prior inflammation, comparable to prior evidence performed with a 28 day recovery period (and despite differences in caerulein protocols). Surprisingly, however, after 3 months, prior inflammation limits ADM in response to secondary injury. With no apparent downregulation of zymogen production or associated tissue damage, this suggests a homeostatic adaptation that does not require metaplasia to resist tissue-level injury. Prior inflammation thus does not serve only to drive lineage plasticity, but to also potentiate or restrain responses in a time-dependent fashion. The molecular mechanisms that allow for acinar cells to ‘tolerate’ a second inflammatory insult warrant further study.

Importantly, inflammatory memory in the pancreas appears to support *Kras-*driven pancreatic tumorigenesis. The iKras model system utilized recently closely recapitulates human PDAC but features overexpression of *Kras*^G12D^, requiring neither injury nor genetically defined tumor suppressor inactivation for progression to malignancy (Ying et al., 2012; Collins et al., 2012). As such, the question of whether inflammation supports the progression of nascent tumors or initial lineage plasticity remains unclear. The model utilized herein not only restricts to the acinar lineage but limits *Kras* expression to endogenous levels and requires inflammation for the loss of acinar cell identity (Guerra et al., 2007; Alonso-Curbelo et al., 2021). Because control animals display only rare PanIN development, we are able to show that it is lineage plasticity that is affected by inflammatory memory and not more facile progression following initial ADM or PanIN formation. Inflammatory memory thus potentiates acquisition of neoplastic cell fate, serving as a substitute for contemporaneous inflammation in systems where the synergy between genetic alteration and environmental insult has been most thoroughly demonstrated (Alonso-Curbelo et al, 2021).

In the nervous system, an essential feature of bona fide memory is recall. Absent memory recall, the findings that we observe could simply represent a persistent change that is unrelated to subsequent cellular responses. Here we demonstrate that differentiated epithelial cells establish memory with *bona fide* ‘recall’ of both epigenetic and transcriptional components of memory. Indeed, the initial response to mutant *Kras* activation in the context of inflammatory memory consists of elevated transcription of specific memory-associated genes. Our single-cell RNA-seq data recapitulate these findings, as *Kras*^*G12D*^ activation in the context of prior inflammation unveils a PanIN cell population not present after *Kras*^G12D^ is turned on in inflammation-naïve mice. In addition to this recall of transcriptional memory, there is also evidence for recall of ‘retained’ epigenetic memory. In particular, putative enhancers and genes identified in our chromatin analyses that are related to and downstream of MAPK signaling -- itself critical to both metaplastic and regenerative components of the response to inflammation (Halbrook et al., 2017) -- are enriched in memory and recall. Thus we show in the pancreas that memory is recovered in a manner that actually impacts the phenotypic outcomes associated with the presence of inflammatory memory (Larsen et al., 2021).

Taken together, our data demonstrate that transcriptional and epigenetic reprogramming in response to inflammation yields persistent alterations to the pancreatic acinar cell; in turn, those precise inflammation-induced alterations are amplified with secondary and substantially delayed oncogenic stress. These data thus demonstrate that long-lived differentiated epithelial cells have the capacity to ‘remember’, and show that the threshold for lineage plasticity can also be durably altered. How this memory is propagated and its specific encoding in chromatin beyond accessibility remain open questions. We and others have demonstrated roles for AP-1 components in defining ‘memory’ chromatin (Naik et al., 2017; Ordovas-Montanes et al., 2018; Lau et al., 2018; de Laval et al., 2020), but closer analyses of TF engagement or histone modifications at key regulatory elements that define memory warrant further study. Specifically, AP-1 has been recently proposed as a universal mediator of memory (Larsen et al., 2021), but the cognate TFs that confer context-specificity to each lineage / response pair are still to be elucidated. This information can in turn guide studies that define whether epigenetic memory can be reversed – by either interfering with the priming or remodeling of enhancers and associated genes or disrupting their recall by secondary stimuli. Both the MAPK pathway and AP-1 transcription factors could represent intriguing targets in this vein. Finally, how the cell-intrinsic memory affects crosstalk with other cell types remains of importance. We do not observe significant changes to either immune or stromal compartments, but how these cell types retain their own memories of inflammation and interact with epithelial cells will be of interest moving forward. Overall, our findings identify inflammatory memory as driving lineage plasticity in early neoplastic cell fate decisions, such that inducing epigenetic ‘amnesia’ of an inflammatory insult could be leveraged as a novel cancer prevention strategy.

## ACKNOWLEDGMENTS

We thank L. Dow, S. Josefowicz, R. Niec, E. Piskounova, J. Pitarresi, and A. Rustgi for critical readings of the manuscript. This work was supported by an American Association for Cancer Research-Pancreatic Cancer Action Network Pathway to Leadership Award (RC), Emerson Collective Cancer Research Fund (RC), and an American Surgical Association Fellowship Award (RC). DJF is supported by a NIH/NCI Ruth L. Kirschstein NRSA F31CA265166; AG is supported by the Prevent Cancer Foundation; AFR is supported by a NCI T32CA203702 grant.

## AUTHOR CONTRIBUTIONS

DJF designed and performed experiments, analyzed data, and wrote the paper. AG performed experiments, analyzed data, and wrote the paper. AO and GP performed experiments. PZ, JLY, AFR, FD, RKY, and JEW analyzed data. OE, DB, and RK supervised data analysis. RC designed, performed, and supervised experiments, analyzed data, supervised data analysis, and wrote the paper.

## STAR METHODS

### Resource availability

#### Lead contact

Further information and requests for resources and reagents should be directed to and will be fulfilled by the lead contact, Rohit Chandwani.

#### Materials availability

This study did not generate new unique reagents.

#### Data and code availability

All high-throughput sequencing data, both raw and processed files, have been deposited in NCBI’s Gene Expression Omnibus and are publicly available as of the date of publication. Accession numbers are listed in the key resources table. Microscopy data reported in this paper will be shared by the lead contact upon request.

This paper does not report original code.

Any additional information required to reanalyze the data reported in this paper is available from the lead contact upon request.

### Experimental model and subject details

#### Animal models

Mice were housed in a pathogen-free facility at Weill Cornell Medicine (WCM). All manipulations were performed under the Institutional Animal Care and Use Committee (IACUC)–approved protocol (2017-0038). Mouse lines used were described previously: Mist1-Cre^ERT2^ (MGI:3821734 Bhlha15^tm3(cre/ERT2)Skz^) (Habbe et al., 2008), LSL-Kras^G12D^ (MGI:2429948 Kras^tm4Tyj^) (Jackson et al., 2001; Tuveson et al., 2004), LSL-tdTomato (MGI:3809523 Gt(ROSA)26Sor^tm9(CAG-tdTomato)Hze^) (Madisen et al., 2010). Male and female animals were used for experiments. Mist1:Cre^ERT2^, LSL-Kras^G12D^, LSL-tdTomato mice were bred to generate Mist1-Cre^ERT2^; LSL-Kras^G12D^; LSL-tdTomato mice, referred to as MKT mice. Mice without Kras^G12D^ are referred to as MT mice.

#### Primary acinar cell culture

Acinar 3D culture were generated as described (Shi et al., 2015) with few modifications. Briefly, acinar cell isolation was obtained with collagenase/dispase mix dissociation as described above, then cells were filtered through a 100 μm cell strainer.

### Method details

#### Tamoxifen treatment

Tamoxifen (Sigma) was dissolved in corn oil and administered by subcutaneous injections (at the indicated ages) at a dosage of 5 mg per injection. Mice were injected once a day for a total of 3 days–– administered every other day (total duration: 5 days). Mice were allowed to recover 1 week after the last Tamoxifen treatment before receiving other treatments (if required).

#### Experimental pancreatitis

For acute pancreatitis treatment, mice received 8 hourly intraperitoneal (IP) injections over two consecutive days of either PBS (saline) or caerulein at a dosage of 50 mg/kg diluted in sterile PBS. Mice were also treated as per a ‘subacute’ pancreatitis protocol with three hourly IP injections a day, three times a week for three weeks or an ‘abbreviated acute’ pancreatitis protocol consisting of three or five hourly injections in a single day, as indicated. For each experiment, pancreata or a portion of pancreata were harvested for histologic analysis.

#### Mouse pancreatic dissociation

Mice were euthanized using CO_2_, and pancreata were excised and placed in 3 mL of ice-cold HBSS while the subsequent pancreata were being harvested. Then, the ice-cold HBSS solution was discarded and pancreata were placed in a 10 cm dish, containing 5 mL of 37°C collagenase and dispase (CD) mix solution–– collagenase D is used for normal tissue; collagenase V is used for fibrotic tissue (CD; ingredients: HBSS w/ Ca2+ Mg2+, Collagenase D / V [1 mg/mL], Dispase II [2 U/mL], STI [0.1 mg/mL], DNase I [0.1 mg/mL]). Pancreata were mechanically dissociated into ∼1-3 mm pieces using razor blades while in CD solution. Minced pancreata were then transferred to 50 mL conical, and an additional 5 mL of CD solution were added, bringing each sample to a final volume of 10 mL. Tubes were placed on a 37°C orbital shaker at 135 rpm for 30 minutes. After incubation, the samples were centrifuged at 1000 rpm for 3 mins, washed with PBS, and then resuspended and incubated with 2 mL of 0.05% Trypsin-EDTA (37°C) for less than 2 min. Trypsin was inactivated immediately after incubation with 10 mL of 37°C PBS/FBS solution (ingredients: PBS w/o Ca2+ Mg2+, FBS [1:5], STI [0.1 mg/mL], DNase I [0.1 mg/mL]). Each tube was then gently inverted twice, centrifuged and resuspended in 10 mL of 37°C FACS buffer (ingredients: PBS, EGTA [10mM], FBS [2%], STI [0.1 mg/mL], DNase I [0.1 mg/mL]). Cell suspension was passed through a 100 μm mesh filter into a new 50 mL conical tube, centrifuged, resuspended with 1 mL of FACS buffer solution with DAPI staining [1:100] and finally transferred to a 40 μm FACS filter tube. All samples were kept on ice before and after FACS. For all FACS, Becton-Dickinson Influx or Becton-Dickinson Aria II were used to collect DAPI-/tdTomato+ cells.

#### Bulk RNA-seq

Following mouse pancreas dissociation, DAPI-/tdTomato+ cells were FACS-sorted directly into TRIzol (100k cells per 750 mL TRIzol) in 1.5 mL microcentrifuge tube. Guanidinium thiocyanate-phenol-chloroform extraction was performed directly and the quality of the samples was determined using the Agilent RNA 6000 Nano Kit on the Agilent Bioanalyzer (Weill Cornell Medicine Genomics Resources Core Facility; WCM-GRCF). The WCM-GRCF prepared libraries using TruSeq RNA Sample Preparation (Non-Stranded and Poly-A selection), and used the NovaSeq 6000 (S1 Flow Cell – Paired End 2×50 cycles) for sequencing. The sequences were aligned to the mouse reference genome (mm9) using STAR4, a universal RNAseq aligner. To improve accuracy of the mapping, the genome was created with a splice junction database based on Gencode vM1 annotation (Harrow et al., 2021). Sequences that mapped to more than one locus were excluded from downstream analysis, since they cannot be confidently assigned. Uniquely mapped sequences were intersected with composite gene models from Gencode vM1 basic annotation using featureCounts (Liao et al., 2014), a tool for assigning sequence reads to genomic features. Composite gene models for each gene consisted of the union of exons of all transcript isoforms of that gene. Uniquely mapped reads that unambiguously overlapped with no more than one Gencode composite gene model were counted for that gene model; the remaining reads were discarded. The counts for each gene model correspond to gene expression values, and were used for subsequent analyses. Prior to the detection of differentially expressed genes, the quality of the sequences was assessed based on several metrics using FastQC and QoRTs (Wingett and Andrews, 2018; Hartley and Mullikin, 2015). Differential gene expression analysis was performed for each comparison using limma voom with default parameters (Law et al., 2014)

#### Bulk ATAC-seq

Following pancreas cell isolation, 50k DAPI-/tdTomato+ acinar cells were sorted into FACS buffer supplemented with FBS [1:10] (Buenrostro et al., 2013). Briefly, acinar cells were centrifuged at 500g for 5 minutes (4°C) and washed with 50 mL of cold PBS. Cells were subsequently lysed using cold lysis buffer and immediately spun at 500 ng for 10 min (4°C). Pellet was resuspended in the transposase reaction mix for the Tn5 tagmentation step for 30 minutes at 37°C and sample was purified using a Qiagen MinElute PCR Purification Kit. Next, DNA was indexed and amplified using PCR. The quality of the samples was assessed using Agilent High Sensitivity DNA kit. DNA libraries were then multiplexed and sequenced on a NextSeq2000 (Paired End; P2 - 100 cycles). For data analysis, quality and adapter filtering was applied to raw reads using ‘trim_galore’ before aligning to mouse assembly mm9 with bowtie2 using the default parameters. The Picard tool MarkDuplicates (http://broadinstitute.github.io/picard/) was used to remove reads with the same start site and orientation. The BEDTools suite (http://bedtools.readthedocs.io) was used to create read density profiles. Enriched regions were discovered using MACS2 and scored against matched input libraries (fold change > 2 and FDR-adjusted p-value < 0.1). A consensus peak atlas was then created by filtering out blacklisted regions (http://mitra.stanford.edu/kundaje/akundaje/release/blacklists/mm9-mouse/mm9-blacklist.bed.gz) and then merging all peaks within 500 bp. A raw count matrix was computed over this atlas using featureCounts (http://subread.sourceforge.net/) with the ‘-p’ option for fragment counting. The count matrix and all genome browser tracks were normalized to a sequencing depth of ten million mapped fragments. DESeq2 was used to classify differential peaks between two conditions using fold change > 2 and FDR-adjusted p-value < 0.1. Peak-gene associations were made using linear genomic distance to the nearest transcription start site with Homer (http://homer.ucsd.edu).

#### Acinar 3D cell culture preparation

Following isolation of acinar cells, cell pellets were resuspended in acinar-explant (AE) media composed of DMEM supplemented with 1% FBS, 1% PS, 1% STI and 1mg/ml of Dexamethasone and then mixed with Matrigel (Corning) in a 2:1 ratio cells:Matrigel. Per 24-well plate, 400 μl of the cells:Matrigel suspension were plated and incubated at 37 °C for solidification for at least 1 hour. Upon Matrigel solidification, 400 μl of warm AE media was added with or without TGF-α at 50ng/ml concentration. Media (with and without TGF-α) was changed at day 1 and day 3 of culture. Brightfield image at 4X and 10X magnification were taken using a Nikon ECLIPSE Ti inverted microscope system equipped with an Andor Zyla 5.5 sCMOS camera.

#### Analysis of brightfield images of spheroids with computer vision

Images were acquired in OME-TIFF format with 16-bit depth in grayscale. Illumination was normalized by subtracting the image passed by a gaussian filter of sigma = 5 * max(x_dim, y_dim) / 300, where x/y_dim are the image dimensions. Images were then inverted and intensity values clipped to the intensity between 3 and 99.8 percentiles, scaling image intensity values to unit range. To segment the spheroid objects from the background, we detected edges in the image by Sobel filtering, and seeded the watershed algorithm using values below 0.3 as background and above 0.95 as positive. We dilated objects in this binary mask with a disk of 3 pixels of diameter, and filtered objects smaller than 32 pixels in diameter. We removed further smaller objects by applying closing (dilation followed by erosion), filling holes within objects, and applying erosion and dilation again, all with a 3 pixel diameter disk. All operations were performed using scikit-image version 0.18.2.11 We then quantified various features for each object in each image using the skimage.measure.regionprops function, and reduced values per image using the mean. Statistical testing was performed between groups of interest with a two-sided Mann–Whitney U test, and adjusted for multiple comparisons with the Benjamini– Hochberg False Discovery Rate method using pingouin (version 0.4.0) (Vallat, 2018).

#### Histopathology and immunofluorescence staining

Pancreata were fixed overnight in 4% buffered PFA, transferred to 70% ethanol, and then embedded in paraffin using IDEXX BioAnalytics laboratory. Serial sections were cut and hematoxylin and eosin (H&E) staining performed. For IF, slides were deparaffinized, underwent an antigen retrieval using Sodium citrate buffer, blocked with 5% BSA supplemented with 0.4% Triton X-100 in PBS, and primary antibodies were incubated overnight at 4°C. Secondary antibodies conjugated with Alexa-488 or Alexa-647 (Invitrogen) were used and DAPI nuclear counterstaining was performed. Fluorescent images were captured with a Nikon ECLIPSE Ti inverted microscope system equipped with an Andor Zyla 5.5 sCMOS camera. For Alcian blue staining, we used the Alcian Blue (ph2.5) Stain Kit (#H-3501) following the manufacturer’s recommendation. H&E and Alcian blue images were captured with a ZEISS Axio Scope.A1 equipped with a Axiocam 105 color.

#### Flow cytometry preparation and analysis

A single cell suspension was generated, and red blood cells (RBCs) removed by using the ACK RBC lysis buffer. Cell suspensions were then stained for extracellular markers (after live/dead staining with Zombie Aqua) per the manufacturer’s instructions, using primary antibodies conjugated to fluorophores in FACS buffer containing 2% FBS. After washing, cells were resuspended in FACS buffer (2% FBS), and analyzed on an Attune NxT flow cytometer. Data analyses were performed with FlowJo software.

#### Single-cell RNA-sequencing

A single cell suspension was generated and subjected to 10x Chromium scRNA-seq as per the manufacturer’s instruction. Analysis was performed using Seurat methods (Stuart et al., 2019). Briefly, raw sequencing data files were preprocessed (e.g. demultiplexed cellular barcodes, read alignment, and generation of gene count matrix) using the Cell Ranger Single Cell Software Suite provided by 10x Genomics. The feature matrix generated by Cell Ranger was used to perform downstream analysis using R toolkit Seurat. Quality control filters were used to exclude low-quality cells. The first filter assessed the level of mitochondrial RNA content in each cell––cells with <7.5% mitochondrial RNA passed the filter. Then, the presence of outliers (e.g. doublets and low detection cells) were determined by evaluating the distribution of cells given the levels of unique RNA transcript number and total RNA transcript numbers––aberrantly high gene count (doublets/multiplets); low gene diversity (low-quality cells/empty droplets). Parameters were set to include cells with gene number values below 4000 and above 800. A filtered Seurat object was subsequently created with high confidence cells that have passed the threshold tests implemented above. Seurat was also implemented for subsequent downstream analyses. To generate a 2-dimensional plot visualizing the different single-cell clusters, default parameters were set for the Uniform Manifold Approximation and Projection (UMAP) method.

### Gene set enrichment analysis

Using normalized read counts of RNA-seq data, the fgseaMultilevel function from the fgsea package with the following parameters were used: minGSSize = 15; maxGSSize = 500 MSigDB Hallmark, MSigDB KEGG, MSigDB REACTOME, and MSigDB Biocarta gene sets were tested for all analyses. Genes were ranked based on DESeq’s wald statistic (stat), which takes into account the log-fold change and its standard error.

### Quantification and statistical analysis

The number of animals is based on feasibility considerations and our interest in observing a relatively large effect size. In addition, a power analysis was performed with a desired power of 0.8, α= 0.05, and a literature search was performed to find expected averages and standard deviations based on similar protocols. Tests for differences between two groups were performed using two-tailed unpaired Student t test or two-tailed Mann–Whitney test as specified in the figure legends. P values were considered significant if less than 0.05. All graphs depict mean ± SEM unless otherwise indicated. Asterisks used to indicate significance correspond with *, P < 0.05; **, P < 0.01; ***, P < 0.001; ****, P < 0.0001. GraphPad Prism 9 (GraphPad Software) was used for statistical analysis of experiments, data processing, and presentation.

