## Supplemental Figures & Figure Legends for "An epigenetic memory of inflammation controls context-dependent lineage plasticity in the pancreas"

#### SUPPLEMENTARY FIGURE LEGENDS

**Figure S1. A,** Gene set enrichment analysis (GSEA) of RNA-seq data performed on tdTomato(+) acinar cells exposed to caerulein and 3 weeks of recovery versus tdTomato(+) acinar cells exposed to caerulein and 2 days of recovery. Top up- and down-regulated pathways are shown. **B,** PCA plot generated of the ATAC-seq data collected from tdTomato(+) acinar cells exposed to either saline or caerulein and allowed to recover for either 2 days or 3 weeks. Each point represents data obtained from a biological replicate mouse. **C,** Heatmap of ChromVar results characterizing the enrichment of known and *de novo* sequence transcription factor motifs in respective conditions. The top 75 motifs are shown. **D,** Venn diagram of motifs upregulated (green) and downregulated (red) in the corresponding contrasts.

**Figure S2. A,** Immunofluorescence for Sox9, TdTomato and DAPI staining of pancreas sections collected 2 days and 12 weeks after caerulein-induced pancreatitis. Representative images shown are from a total of N=2-5 mice per condition. **B,** Immunofluorescence for Cpa1, CK19 and DAPI staining of pancreas sections collected 2 days and 12 weeks after caerulein-induced pancreatitis. Representative images shown are from a total of N=2-5 mice per condition. **C,** UMAP projection of single-cell RNA-seq data generated with cells isolated from pancreata collected 2 days and 6 weeks after saline or caerulein treatment. **D,** Feature plots generated using single-cell RNA-seq data (subset on the epithelial compartment) for the indicated acinar/ductal/progenitor transcripts. **E,** Heatmap of single cell RNA-seq data on the epithelial compartment; Top 50 differentially expressed genes identified by comparing ductal versus acinar cell clusters (log2FC > 1), wherein each column represents a single cell. **F,** Heatmap of all differentially expressed genes identified by comparing caerulein + 2d recovery and saline + 2d recovery (padj. < 0.01). **G,** Representative chromatin accessibility tracks at PanIN/duct/acinar genes visualized with IGV; light-blue boxes highlight chromatin dynamics that are up at 2 days and 3 weeks of recovery and lost with prolonged recovery; green boxes denote persistently accessible regions; dark-blue boxes represent regions that are lost and then regained with prolonged recovery. Scale bar for all tracks 0-60.

**Figure S3. A & C,** Schematic representations of primary pancreatitis and inflammatory re-challenge treatment regimen in wild-type mice. **B & D,** Hematoxylin and eosin staining of inflammation-naïve and inflammation-resolved mouse pancreas sections collected 2 days after inflammatory re-challenge. Representative images from a total of N=3-5 mice per condition (B) or images from all mice (D) are shown.

**Figure S4. A,** Schematic representation of lineage-traced mouse model exposed to primary treatment (saline or caerulein), 3 weeks of recovery, and then 3 weeks of mutant Kras activation. **B,** H&E and Alcian blue staining of mouse pancreas sections collected from mice previously exposed to either saline or caerulein plus 3 weeks of recovery, then 3 weeks of $Kras^{G12D}$. Representative images shown are from a total of N=2-5 mice per condition. **C,** H&E staining of inflammation-naïve and inflammation-resolved lineage-traced mice, after 12 weeks of recovery and then 2 days of $Kras^{G12D}$. **D & E,** Immunofluorescence for **(D)** Cpa1 and CK19 and **(E)** Dclk1 and DAPI staining of pancreas sections collected from inflammation-naïve and inflammation-resolved mice after 12 weeks of recovery and then 2 days of mutant Kras. **F,** GSEA of RNA-seq data comparing inflammation-naïve and inflammation-resolved tdTomato(+) acinar cells after 12 weeks of recovery and then 2 days of $Kras^{G12D}$. **G,** Representative chromatin accessibility tracks at PanIN-specific genes visualized with IGV; light green boxes highlight persistently accessible regions.

### Figure S1

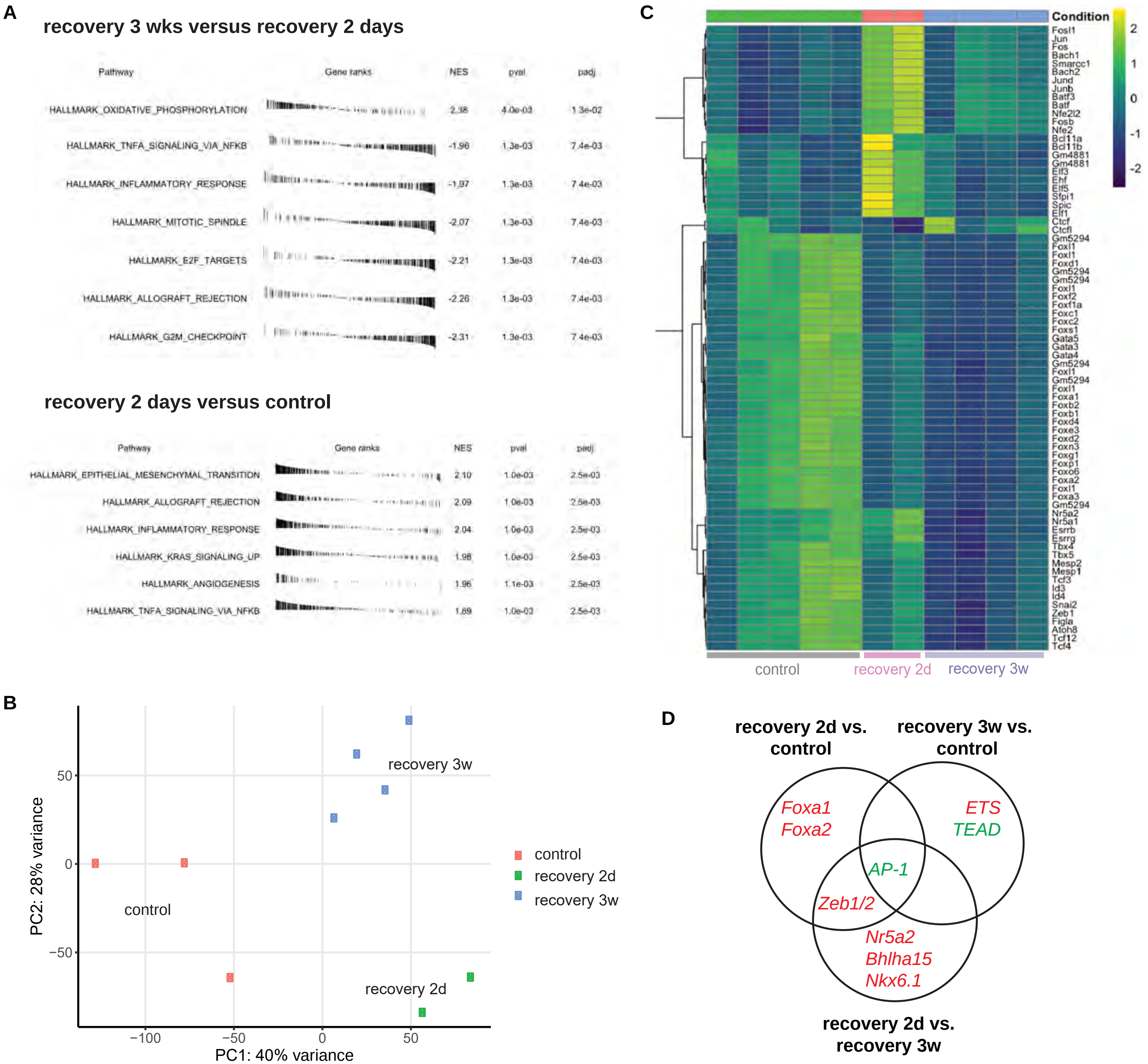
A
recovery 3 wks versus recovery 2 days
Pathway Gene ranks NES pval padj
HALLMARK_OXIDATIVE_PHOSPHORYLATION 2.38 4.0e-03 1.3e-02
HALLMARK_TNFA_SIGNALING_VIA_NFKB -1.96 1.3e-03 7.4e-03
HALLMARK_INFLAMMATORY_RESPONSE -1.97 1.3e-03 7.4e-03
HALLMARK_MITOTIC_SPINDLE -2.07 1.3e-03 7.4e-03
HALLMARK_E2F_TARGETS -2.21 1.3e-03 7.4e-03
HALLMARK_ALLOGRAFT_REJECTION -2.26 1.3e-03 7.4e-03
HALLMARK_G2M_CHECKPOINT -2.31 1.3e-03 7.4e-03
recovery 2 days versus control
Pathway Gene ranks NES pval padj
HALLMARK_EPITHELIAL_MESENCHYMAL_TRANSITION 2.10 1.0e-03 2.5e-03
HALLMARK_ALLOGRAFT_REJECTION 2.09 1.0e-03 2.5e-03
HALLMARK_INFLAMMATORY_RESPONSE 2.04 1.0e-03 2.5e-03
HALLMARK_KRAS_SIGNALING_UP 1.98 1.0e-03 2.5e-03
HALLMARK_ANGIOGENESIS 1.96 1.1e-03 2.5e-03
HALLMARK_TNFA_SIGNALING_VIA_NFKB 1.89 1.0e-03 2.5e-03
B
PC2: 28% variance
PC1: 40% variance
50
0
−50
−100
−50
0
50
recovery 3w
control
recovery 2d
control
recovery 2d
recovery 3w
C
Condition
2
1
0
-1
-2
Fosl1 Jun Fos Bach1 Smarcc1 Bach2 Jund Junb Batf3 Batf Nfe2l2 Fosb Nfe2 Bcl11a Bcl11b Gm4881 Gm4881 Elf3 Ehf Elf5 Sfpi1 Spic Elf1 Ctcf Ctcfl Gm5294 Foxl1 Foxl1 Foxd1 Gm5294 Gm5294 Foxl1 Foxf2 Foxf1a Foxc1 Foxc2 Foxs1 Gata5 Gata3 Gata4 Gm5294 Foxl1 Gm5294 Foxl1 Foxa1 Foxb2 Foxb1 Foxd4 Foxe3 Foxd2 Foxn3 Foxg1 Foxp1 Foxo6 Foxa2 Foxl1 Foxa3 Gm5294 Nr5a2 Nr5a1 Esrrb Esrrg Tbx4 Tbx5 Mesp2 Mesp1 Tcf3 Id3 Id4 Snai2 Zeb1 Figla Atoh8 Tcf12 Tcf4
control
recovery 2d
recovery 3w
D
recovery 2d vs. control
recovery 3w vs. control
recovery 2d vs. recovery 3w
Foxa1
Foxa2
ETS
TEAD
AP-1
Zeb1/2
Nr5a2
Bhlha15
Nkx6.1

#### Figure S2

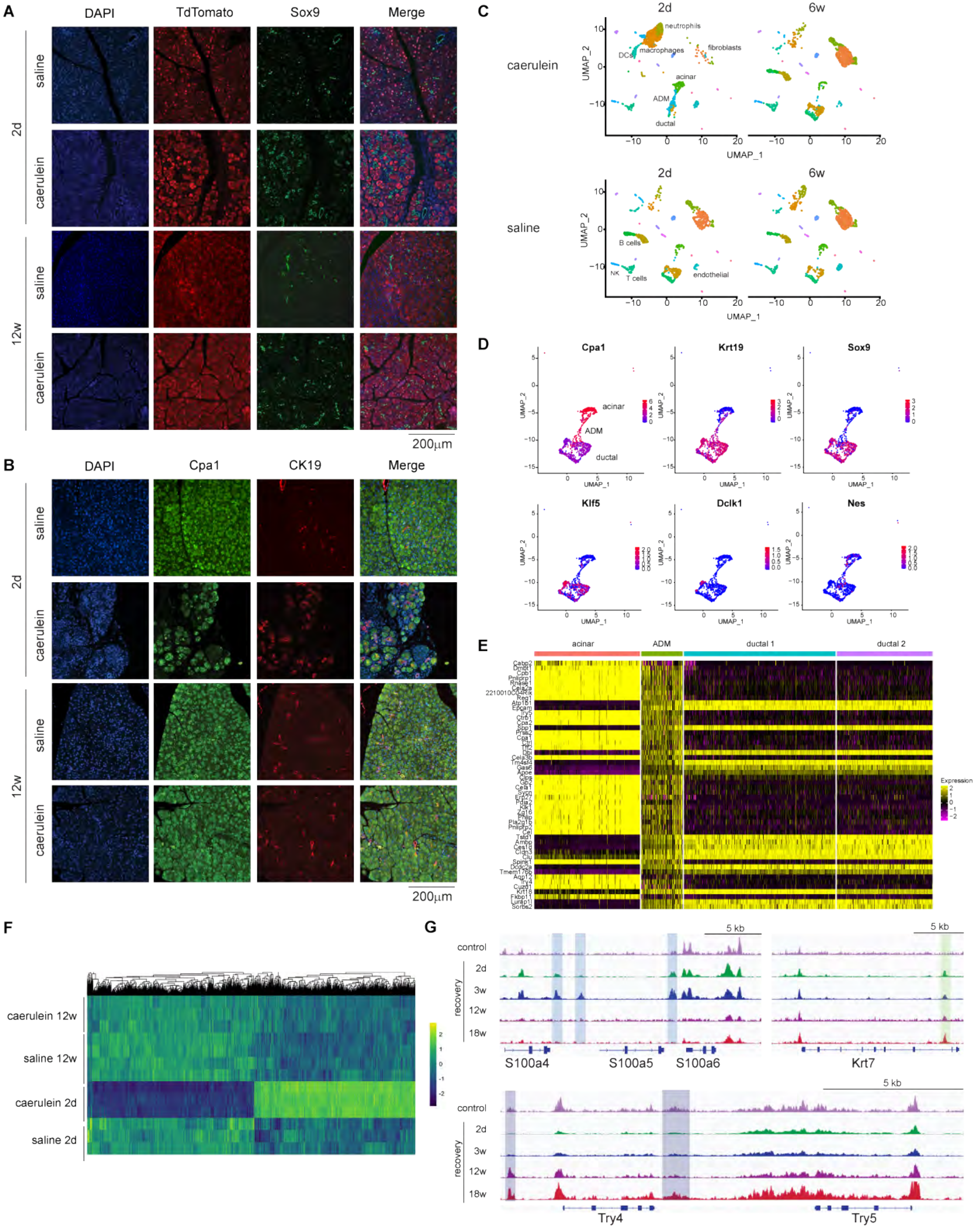
A
DAPI
TdTomato
Sox9
Merge
2d
saline
caerulein
12w
saline
caerulein
200μm
B
DAPI
Cpa1
CK19
Merge
2d
saline
caerulein
12w
saline
caerulein
200μm
C
2d
6w
caerulein
saline
neutrophils
macrophages
fibroblasts
DCs
acinar
ADM
ductal
B cells
NK
T cells
endothelial
UMAP_1
UMAP_2
D
Cpa1
Krt19
Sox9
Klf5
Dclk1
Nes
acinar
ADM
ductal
E
acinar
ADM
ductal 1
ductal 2
Expression
F
caerulein 12w
saline 12w
caerulein 2d
saline 2d
G
control
recovery
2d
3w
12w
18w
5 kb
S100a4
S100a5
S100a6
Krt7
Try4
Try5

#### Figure S3

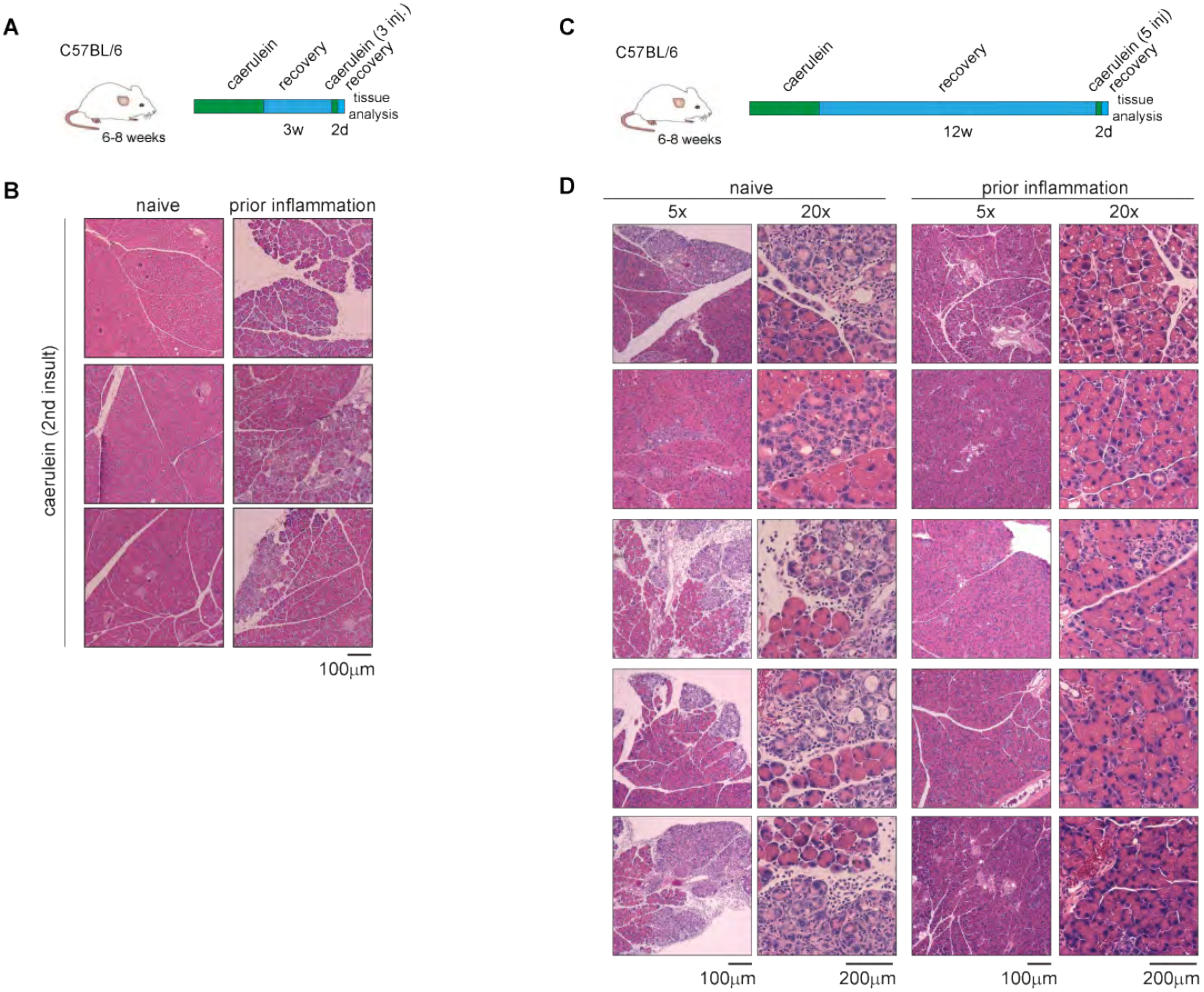
A
C57BL/6
6-8 weeks
caerulein
recovery
caerulein (3 inj.)
recovery
tissue analysis
3w
2d
B
naive
prior inflammation
caerulein (2nd insult)
100μm
C
C57BL/6
6-8 weeks
caerulein
recovery
caerulein (5 inj)
recovery
tissue analysis
12w
2d
D
naive
prior inflammation
5x
20x
5x
20x
100μm
200μm
100μm
200μm

### Figure S4

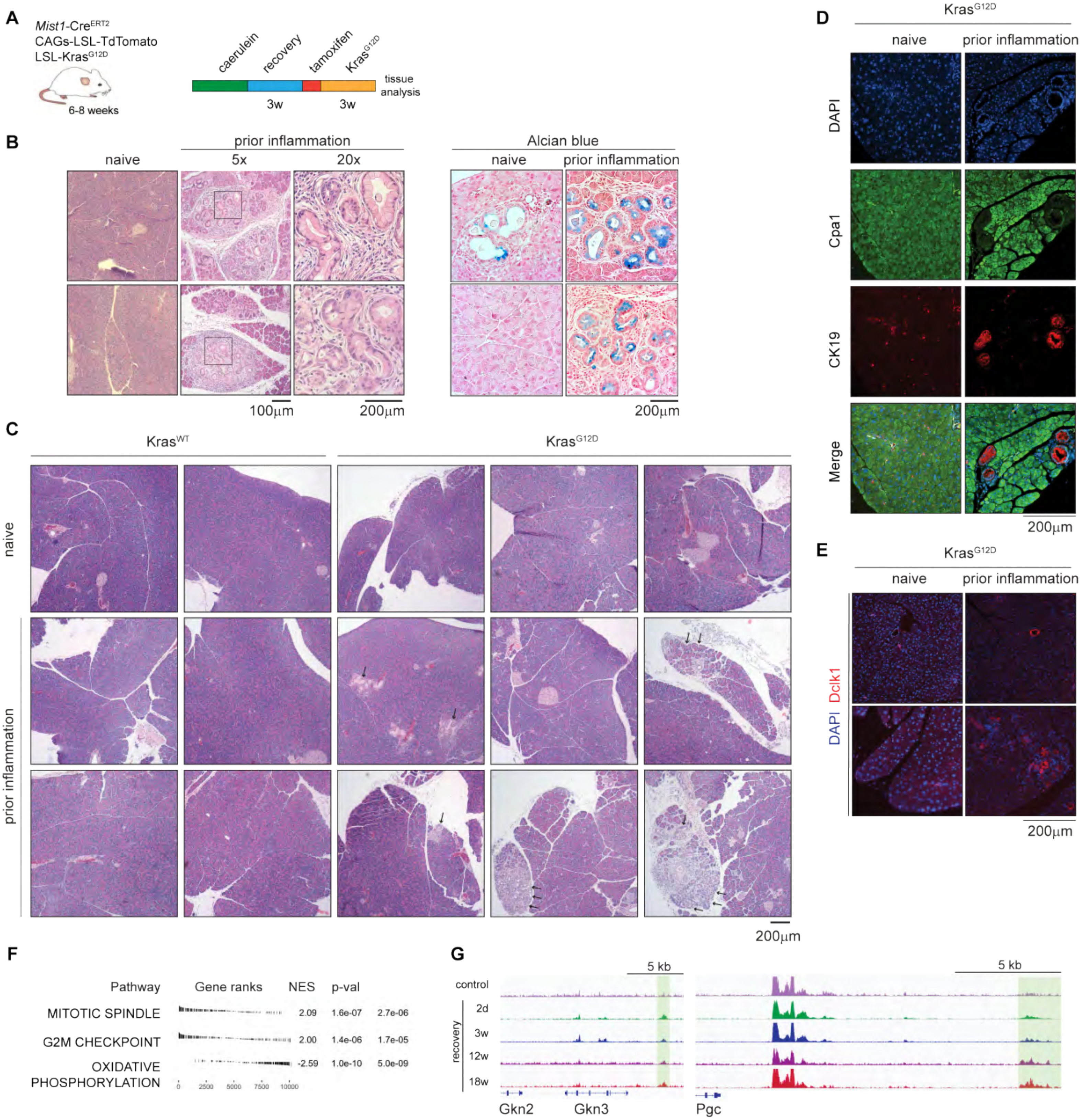
A
Mist1-CreERT2
CAGs-LSL-TdTomato
LSL-KrasG12D
6-8 weeks
caerulein
recovery
tamoxifen
KrasG12D
tissue analysis
3w
3w
B
naive
prior inflammation
5x
20x
Alcian blue
naive
prior inflammation
100µm
200µm
200µm
C
KrasWT
KrasG12D
naive
prior inflammation
200µm
D
KrasG12D
naive
prior inflammation
DAPI
Cpa1
CK19
Merge
200µm
E
KrasG12D
naive
prior inflammation
DAPI Dclk1
200µm
F
Pathway
Gene ranks
NES
p-val
MITOTIC SPINDLE 2.09 1.6e-07 2.7e-06
G2M CHECKPOINT 2.00 1.4e-06 1.7e-05
OXIDATIVE PHOSPHORYLATION -2.59 1.0e-10 5.0e-09
0 2500 5000 7500 10000
G
5 kb
5 kb
control
recovery
2d
3w
12w
18w
Gkn2
Gkn3
Pgc
